## Supplementary_Figures for "Mutation density changes in SARS-CoV-2 are related to the pandemic stage but to a lesser extent in the dominant strain with mutations in spike and RdRp"

**Figure S1**

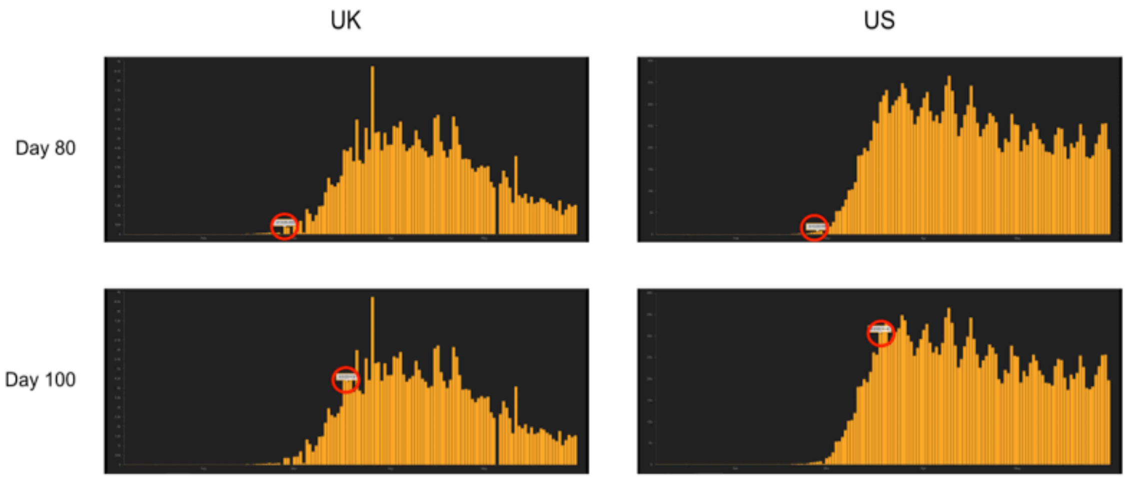


**Figure S1.** Barplot of daily case numbers for UK and US. Red circles indicate day 80 for the top two plots, and day 100 for the bottom two plots. Plots are generated using COVID-19 Dashboard by the Center for Systems Science and Engineering (CSSE) at Johns Hopkins University.

#### Figure S2

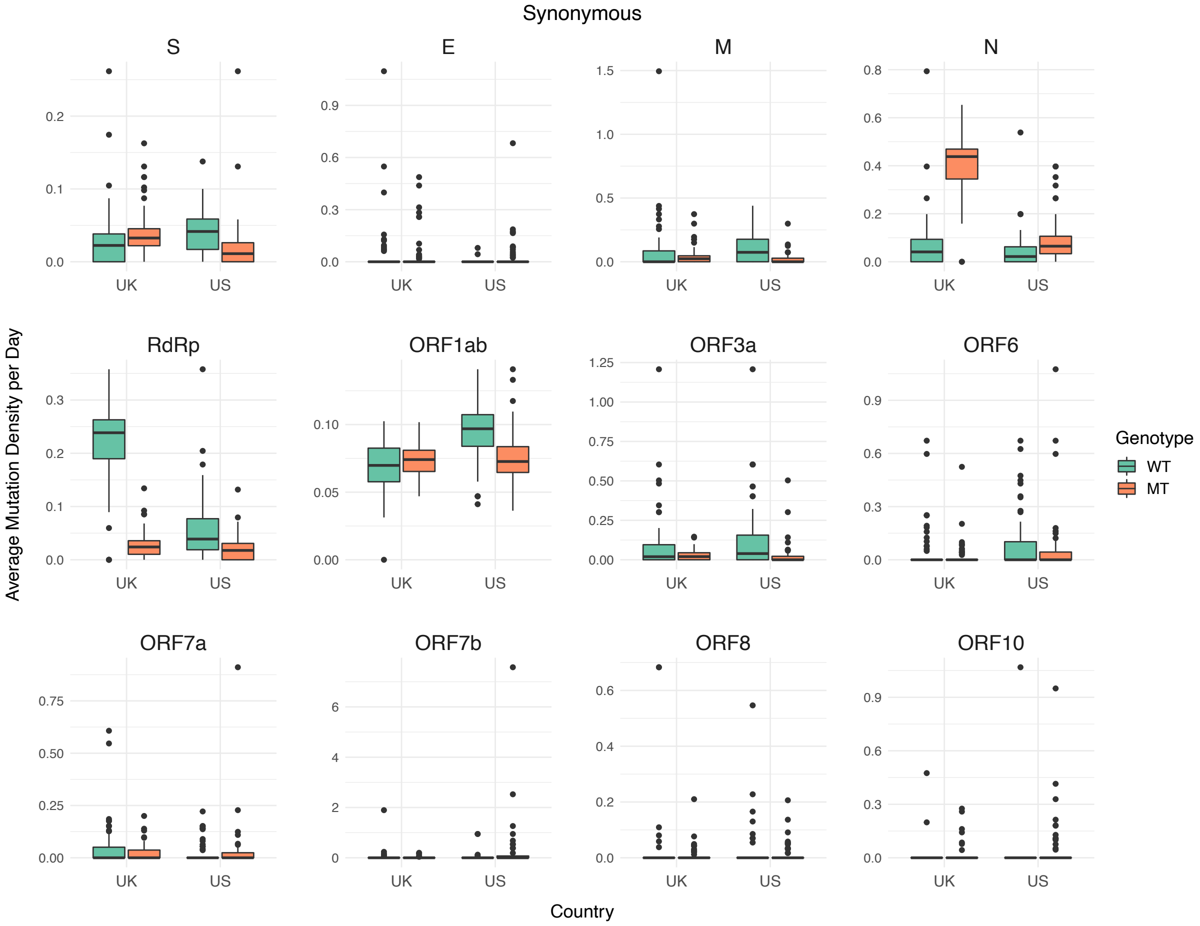


**Figure S2.** Boxplot of synonymous mutation densities in WT and MT isolates, divided by coding region. Horizontal line in box represents median value, the bottom and top whiskers represent the lower and upper quartile, and the dots represent outlier values. Green boxes correspond to WT isolates, while orange boxes correspond to MT isolates.

### Figure S3

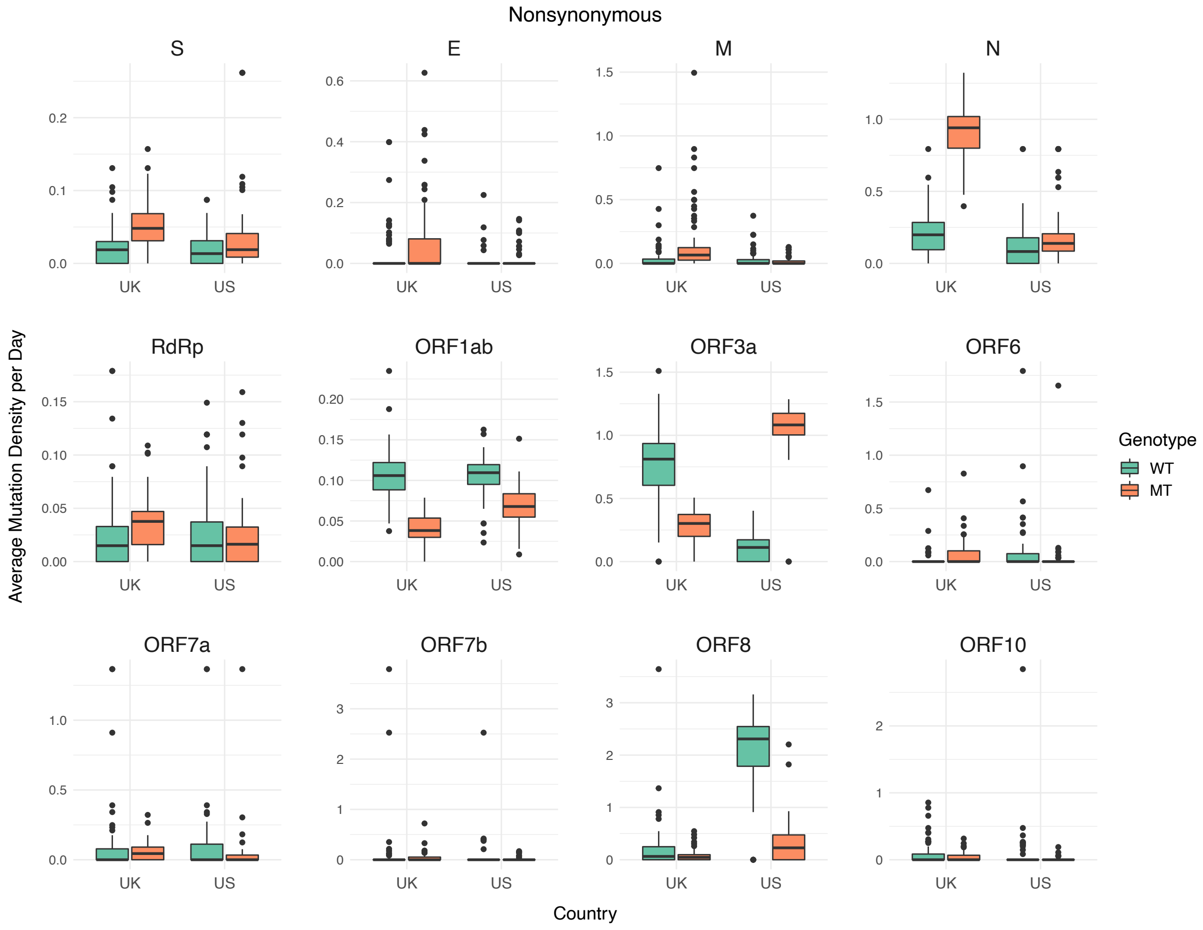


**Figure S3.** Boxplot of nonsynonymous mutation densities in WT and MT isolates, divided by coding region. Horizontal line in box represents median value, the bottom and top whiskers represent the lower and upper quartile, and the dots represent outlier values. Green boxes correspond to WT isolates, while orange boxes correspond to MT isolates.

### Figure S4

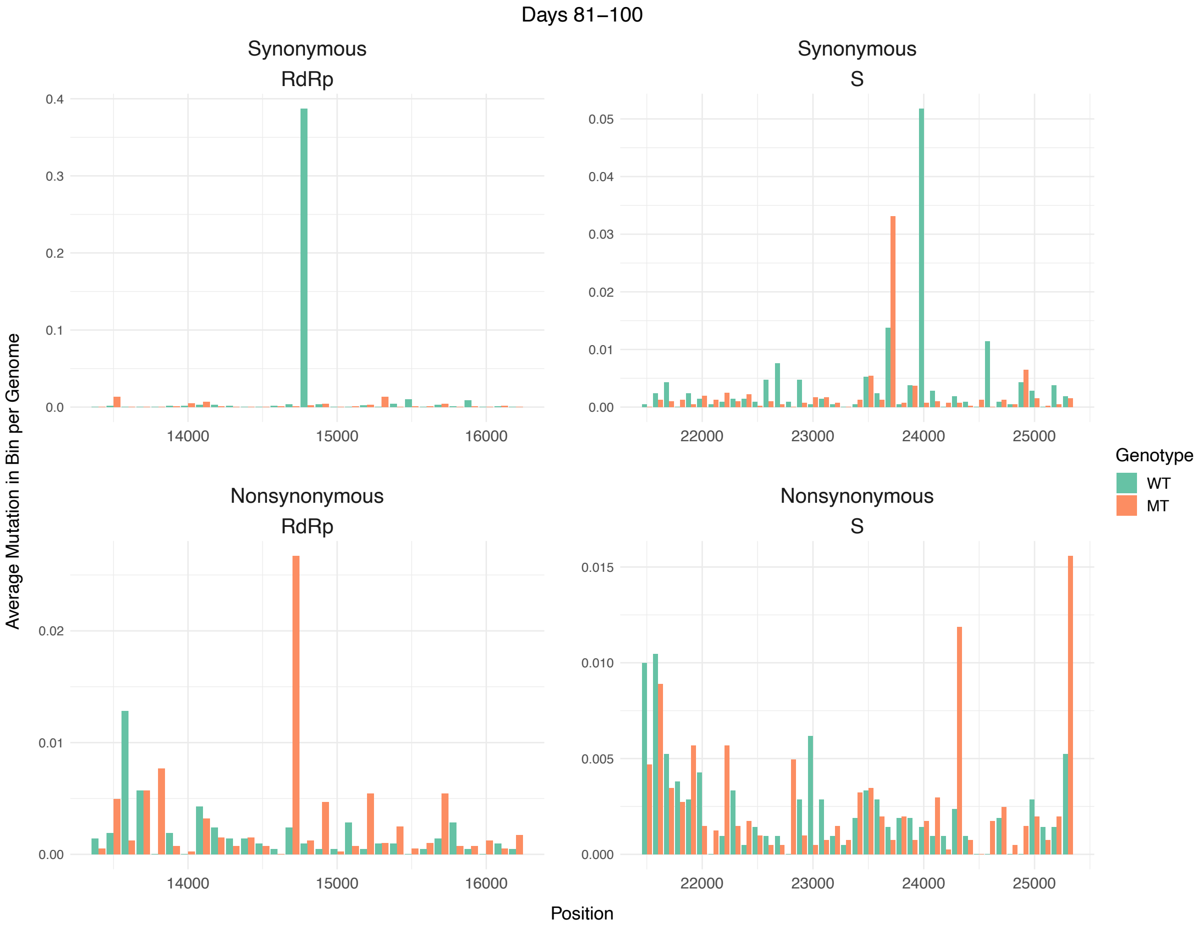


**Figure S4.** Distribution of mutations across RdRp and S coding regions in WT and MT samples between days 81 and 100. Each bar represents the total of all mutations in a 100-nucleotide long region divided by number of isolates in the indicated sample type in the time period, with green bars indicating WT isolates and orange bars indicating MT isolates.

#### Figure S5

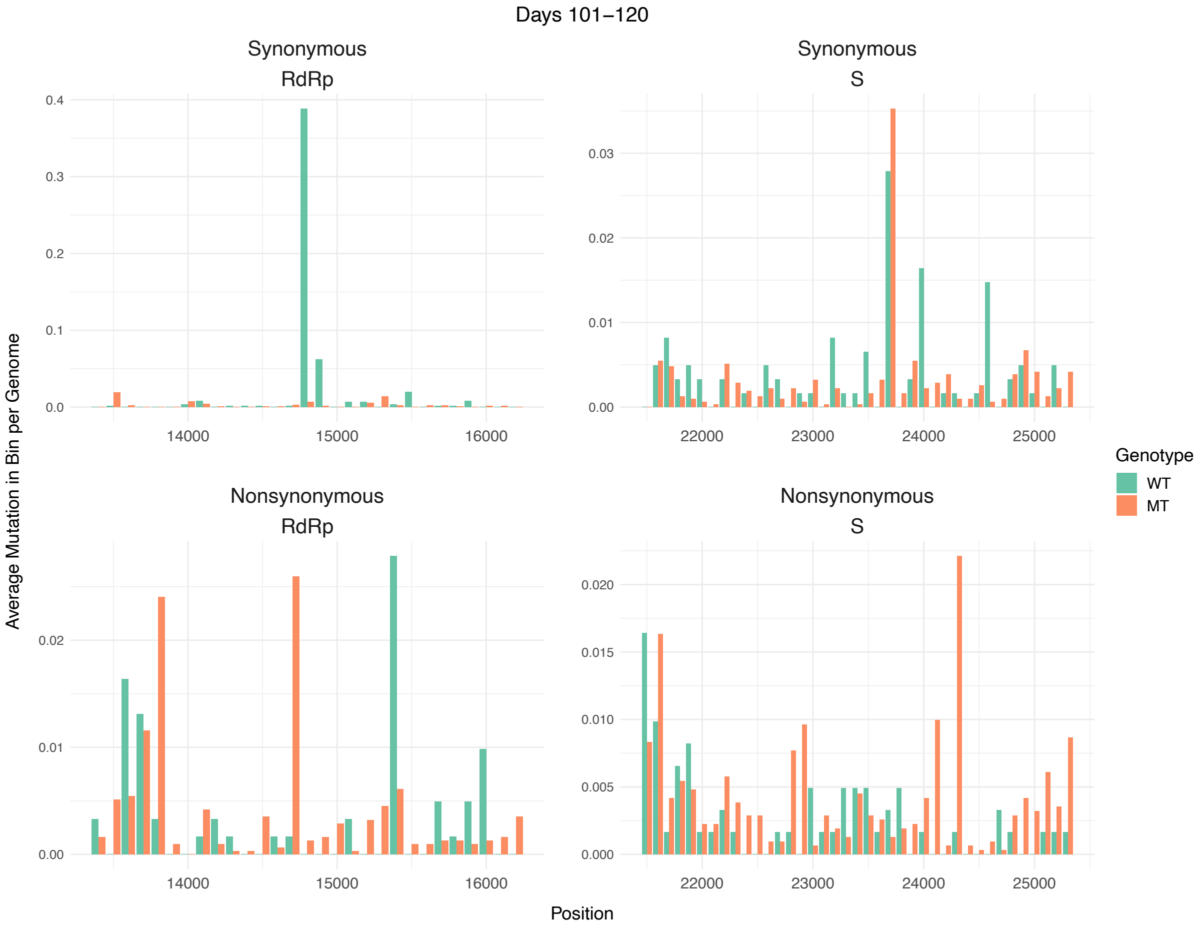


**Figure S5.** Distribution of mutations across RdRp and S coding regions in WT and MT samples between days 101 and 120. Each bar represents the total of all mutations in a 100-nucleotide long region divided by number of isolates in the indicated sample type in the time period, with green bars indicating WT isolates and orange bars indicating MT isolates.
