## Supplementary_Tables for "Mutation density changes in SARS-CoV-2 are related to the pandemic stage but to a lesser extent in the dominant strain with mutations in spike and RdRp"

**Supplementary Table 1.** Comparisons of the wild type and mutant groups in UK and US isolates for each SARS-CoV-2 gene.

| **Variable** | **Country** | **Median (IQR)**  **(WT vs. MT)** | **Mean Rank**  **(WT vs. MT)** | **p*** |
| --- | --- | --- | --- | --- |
| S synonymous | UK | 0.02 (0.03) vs. 0.03 (0.02) | 61.13 vs. 79.93 | 0.006* |
|  | US | 0.04 (0.04) vs. 0.01 (0.02) | 81.17 vs. 51.51 | <0.001* |
| S nonsynonymous | UK | 0.01 (0.03) vs. 0.05 (0.04) | 48.88 vs. 91.03 | <0.001* |
|  | US | 0.01 (0.03) vs. 0.02 (0.03) | 58.95 vs. 72.73 | 0.035* |
| N synonymous | UK | 0.04 (0.09) vs. 0.43 (0.18) | 37.21 vs. 101.59 | <0.001* |
|  | US | 0.02 (0.06) vs. 0.06 (0.08) | 54.05 vs. 77.41 | <0.001* |
| N nonsynonymous | UK | 0.19 (0.23) vs. 0.94 (0.28) | 34.45 vs. 104.09 | <0.001* |
|  | US | 0.08 (0.18) vs. 0.14 (0.12) | 55.52 vs. 76.01 | 0.002* |
| E synonymous | UK | 0.00 (0.00) vs. 0.00 (0.00) | 70.54 vs. 71.42 | 0.852 |
|  | US | 0.00 (0.00) vs. 0.00 (0.00) | 59.48 vs. 72.22 | 0.001* |
| E nonsynonymous | UK | 0.00 (0.00) vs. 0.00 (0.09) | 62.84 vs. 78.39 | 0.008* |
|  | US | 0.00 (0.00) vs. 0.00 (0.00) | 63.20 vs. 68.67 | 0.147 |
| M synonymous | UK | 0.00 (0.00) vs. 0.02 (0.05) | 69.76 vs. 72.12 | 0.715 |
|  | US | 0.07 (0.18) vs. 0.00 (0.03) | 79.05 vs. 53.53 | <0.001* |
| M nonsynonymous | UK | 0.00 (0.03) vs. 0.07 (0.10) | 51.20 vs. 88.93 | <0.001* |
|  | US | 0.00 (0.04) vs. 0.00 (0.02) | 65.86 vs. 66.13 | 0.961 |
| RdRp synonymous | UK | 0.24 (0.08) vs. 0.02 (0.03) | 104.91 vs. 40.30 | <0.001* |
|  | US | 0.04 (0.06) vs. 0.02 (0.03) | 80.26 vs. 52.38 | <0.001* |
| RdRp nonsynonymous | UK | 0.02 (0.03) vs. 0.04 (0.03) | 58.57 vs. 82.25 | <0.001* |
|  | US | 0.02 (0.04) vs. 0.02 (0.03) | 65.50 vs. 66.48 | 0.881 |
| Orf1ab synonymous | UK | 0.07 (0.03) vs. 0.07 (0.02) | 65.48 vs. 76.00 | 0.126 |
|  | US | 0.10 (0.02) vs. 0.07 (0.02) | 83.36 vs. 49.42 | <0.001* |
| Orf1ab nonsynonymous | UK | 0.11 (0.04) vs. 0.03 (0.02) | 105.51 vs. 39.76 | <0.001* |
|  | US | 0.11 (0.03) vs. 0.07 (0.03) | 91.35 vs. 41.78 | <0.001* |
| Orf3a synonymous | UK | 0.02 (0.10) vs. 0.02 (0.05) | 74.29 vs. 68.02 | 0.341 |
|  | US | 0.04 (0.17) vs. 0.00 (0.02) | 76.95 vs. 55.54 | 0.001* |
| Orf3a nonsynonymous | UK | 0.81 (0.33) vs. 0.30 (0.18) | 102.19 vs. 42.76 | <0.001* |
|  | US | 0.11 (0.17) vs. 1.08 (0.18) | 35.05 vs. 95.56 | <0.001* |
| Orf6 synonymous | UK | 0.00 (0.00) vs. 0.00 (0.00) | 71.12 vs. 70.89 | 0.963 |
|  | US | 0.00 (0.11) vs. 0.00 (0.05) | 66.94 vs. 65.10 | 0.725 |
| Orf6 nonsynonymous | UK | 0.00 (0.00) vs. 0.00 (0.10) | 59.05 vs. 81.82 | <0.001* |
|  | US | 0.00 (0.12) vs. 0.00 (0.00) | 71.27 vs. 60.97 | 0.026* |
| Orf7a synonymous | UK | 0.00 (0.05) vs. 0.00 (0.04) | 74.08 vs. 68.21 | 0.316 |
|  | US | 0.00 (0.00) vs. 0.00 (0.03) | 64.28 vs. 67.64 | 0.502 |
| Orf7a nonsynonymous | UK | 0.00 (0.08) vs. 0.04 (0.09) | 64.51 vs. 76.88 | 0.051 |
|  | US | 0.00 (0.12) vs. 0.00 (0.03) | 71.45 vs. 60.79 | 0.066 |
| Orf7b synonymous | UK | 0.00 (0.00) vs. 0.00 (0.00) | 72.94 vs. 69.24 | 0.248 |
|  | US | 0.00 (0.00) vs. 0.00 (0.08) | 57.66 vs. 73.96 | <0.001* |
| Orf7b nonsynonymous | UK | 0.00 (0.00) vs. 0.00 (0.06) | 64.63 vs. 76.76 | 0.015* |
|  | US | 0.00 (0.00) vs. 0.00 (0.00) | 65.97 vs. 66.03 | 0.986 |
| Orf8 synonymous | UK | 0.00 (0.00) vs. 0.00 (0.00) | 68.23 vs. 73.51 | 0.197 |
|  | US | 0.00 (0.00) vs. 0.00 (0.00) | 63.80 vs. 68.10 | 0.320 |
| Orf8 nonsynonymous | UK | 0.06 (0.25) vs. 0.05 (0.10) | 75.26 vs. 67.14 | 0.218 |
|  | US | 2.31 (0.85) vs. 0.23 (0.48) | 95.52 vs. 37.81 | <0.001* |
| Orf10 synonymous | UK | 0.00 (0.00) vs. 0.00 (0.00) | 68.68 vs. 73.10 | 0.130 |
|  | US | 0.00 (0.00) vs. 0.00 (0.00) | 59.63 vs. 72.08 | 0.001* |
| Orf10 nonsynonymous | UK | 0.00 (0.09) vs. 0.00 (0.07) | 70.88 vs. 71.11 | 0.967 |
|  | US | 0.00 (0.00) vs. 0.00 (0.00) | 70.77 vs. 61.45 | 0.019* |

WT, Wild type; MT; Mutant; IQR, Interquartile Range; * Wilcoxon rank-sum (Mann-Whitney) test; *p-value<0.05 is significant.

**Supplementary Table 2.** Comparisons of the days groups in wild type and mutant groups in UK and US isolates for each SARS-CoV-2 gene.

| **Variable** | **Country** | **Phenotype** | **Median (IQR)**  **(60-100 vs. 101-140)** | **Mean Rank**  **(60-100 vs. 101-140)** | **p*** |
| --- | --- | --- | --- | --- | --- |
| S synonymous | UK | WT | 0.02 (0.02) vs. 0.02 (0.05) | 34.17 vs. 33.81 | 0.939 |
|  |  | MT | 0.03 (0.04) vs. 0.04 (0.02) | 30.90 vs. 43.75 | 0.010* |
|  | USA | WT | 0.04 (0.03) vs. 0.03 (0.06) | 36.36 vs. 27.60 | 0.054 |
|  |  | MT | 0.01 (0.02) vs. 0.02 (0.03) | 27.89 vs. 40.69 | 0.006* |
| S nonsynonymous | UK | WT | 0.02 (0.03) vs. 0.03 (0.04) | 33.81 vs. 34.23 | 0.928 |
|  |  | MT | 0.03 (0.04) vs. 0.06 (0.03) | 29.67 vs. 44.92 | 0.002* |
|  | USA | WT | 0.02 (0.02) vs. 0.00 (0.01) | 40.40 vs. 22.97 | <0.001* |
|  |  | MT | 0.02 (0.02) vs. 0.04 (0.06) | 26.50 vs. 42.20 | 0.001* |
| N synonymous | UK | WT | 0.04 (0.06) vs. 0.05 (0.13) | 32.06 vs. 36.26 | 0.362 |
|  |  | MT | 0.41 (0.13) vs. 0.44 (0.12) | 33.13 vs. 41.64 | 0.088 |
|  | USA | WT | 0.04 (0.06) vs. 0.00 (0.04) | 38.01 vs. 25.84 | 0.007* |
|  |  | MT | 0.06 (0.05) vs. 0.07 (0.11) | 33.17 vs. 34.91 | 0.715 |
| N nonsynonymous | UK | WT | 0.16 (0.21) vs. 0.26 (0.19) | 25.61 vs. 43.73 | <0.001* |
|  |  | MT | 0.86 (0.25) vs. 0.98 (0.13) | 28.18 vs. 46.33 | <0.001* |
|  | USA | WT | 0.10 (0.17) vs. 0.06 (0.15) | 35.56 vs. 28.81 | 0.144 |
|  |  | MT | 0.13 (0.10) vs. 0.15 (0.17) | 32.06 vs. 36.13 | 0.393 |
| E synonymous | UK | WT | 0.00 (0.00) vs. 0.00 (0.00) | 34.08 vs. 33.90 | 0.955 |
|  |  | MT | 0.00 (0.00) vs. 0.00 (0.02) | 34.08 vs. 40.74 | 0.058 |
|  | USA | WT | 0.00 (0.00) vs. 0.00 (0.00) | 33.33 vs. 31.50 | 0.194 |
|  |  | MT | 0.00 (0.03) vs. 0.00 (0.00) | 35.41 vs. 32.45 | 0.395 |
| E nonsynonymous | UK | WT | 0.00 (0.05) vs. 0.00 (0.00) | 34.36 vs. 33.58 | 0.823 |
|  |  | MT | 0.02 (0.00) vs. 0.03 (0.00) | 24.68 vs. 49.64 | <0.001* |
|  | USA | WT | 0.07 (0.18) vs. 0.00 (0.00) | 34.57 vs. 30.00 | 0.036* |
|  |  | MT | 0.00 (0.00) vs. 0.00 (0.00) | 33.96 vs. 34.05 | 0.977 |
| M synonymous | UK | WT | 0.00 (0.08) vs. 0.00 (0.10) | 34.18 vs. 33.79 | 0.928 |
|  |  | MT | 0.00 (0.03) vs. 0.04 (0.07) | 31.17 vs. 43.50 | 0.010* |
|  | USA | WT | 0.15 (0.15) vs. 0.00 (0.00) | 43.39 vs. 19.36 | <0.001* |
|  |  | MT | 0.00 (0.03) vs. 0.00 (0.02) | 34.91 vs. 33.00 | 0.640 |
| M nonsynonymous | UK | WT | 0.00 (0.03) vs. 0.00 (0.04) | 35.72 vs. 32.00 | 0.363 |
|  |  | MT | 0.10 (0.41) vs. 0.06 (0.09) | 44.86 vs. 30.53 | 0.004* |
|  | USA | WT | 0.00 (0.04) vs. 0.00 (0.00) | 34.76 vs. 29.78 | 0.202 |
|  |  | MT | 0.01 (0.03) vs. 0.00 (0.00) | 39.44 vs. 28.05 | 0.006* |
| RdRp synonymous | UK | WT | 0.23 (0.07) vs. 0.24 (0.08) | 30.22 vs. 38.39 | 0.087 |
|  |  | MT | 0.01 (0.03) vs. 0.03 (0.03) | 26.94 vs. 47.50 | <0.001* |
|  | USA | WT | 0.04 (0.05) vs. 0.05 (0.13) | 31.96 vs. 33.16 | 0.797 |
|  |  | MT | 0.01 (0.02) vs. 0.03 (0.05) | 29.76 vs. 38.64 | 0.058 |
| RdRp nonsynonymous | UK | WT | 0.02 (0.03) vs. 0.00 (0.04) | 37.75 vs. 33.13 | 0.724 |
|  |  | MT | 0.02 (0.04) vs. 0.04 (0.04) | 28.58 vs. 45.95 | <0.001* |
|  | USA | WT | 0.01 (0.02) vs. 0.02 (0.08) | 30.64 vs. 34.74 | 0.371 |
|  |  | MT | 0.02 (0.03) vs. 0.02 (0.04) | 32.80 vs. 35.31 | 0.594 |
| Orf1ab synonymous | UK | WT | 0.06 (0.02) vs. 0.08 (0.02) | 25.19 vs. 44.24 | <0.001* |
|  |  | MT | 0.07 (0.01) vs. 0.08 (0.01) | 19.39 vs. 54.66 | <0.001* |
|  | USA | WT | 0.10 (0.01) vs. 0.09 (0.05) | 33.74 vs. 31.00 | 0.557 |
|  |  | MT | 0.07 (0.01) vs. 0.08 (0.02) | 23.81 vs. 45.14 | <0.001* |
| Orf1ab nonsynonymous | UK | WT | 0.10 (0.04) vs. 0.11 (0.04) | 27.57 vs. 41.47 | 0.004* |
|  |  | MT | 0.03 (0.02) vs. 0.05 (0.02) | 22.17 vs. 52.03 | <0.001* |
|  | USA | WT | 0.10 (0.02) vs. 0.11 (0.05) | 32.16 vs. 32.91 | 0.871 |
|  |  | MT | 0.06 (0.03) vs. 0.08 (0.03) | 26.39 vs. 42.33 | 0.001* |
| Orf3a synonymous | UK | WT | 0.02 (0.06) vs. 0.03 (0.15) | 31.28 vs. 37.16 | 0.194 |
|  |  | MT | 0.00 (0.03) vs. 0.03 (0.06) | 31.69 vs. 43.00 | 0.019* |
|  | USA | WT | 0.03 (0.09) vs. 0.13 (0.40) | 28.16 vs. 37.74 | 0.033* |
|  |  | MT | 0.00 (0.02) vs. 0.00 (0.03) | 33.90 vs. 34.11 | 0.961 |
| Orf3a nonsynonymous | UK | WT | 0.80 (0.30) vs. 0.89 (0.40) | 31.75 vs. 36.61 | 0.308 |
|  |  | MT | 0.23 (0.33) vs. 0.35 (0.13) | 28.82 vs. 45.72 | 0.001* |
|  | USA | WT | 0.12 (0.09) vs. 0.00 (0.19) | 36.66 vs. 27.48 | 0.047* |
|  |  | MT | 1.05 (0.15) vs. 1.13 (0.17) | 29.84 vs. 38.55 | 0.067 |
| Orf6 synonymous | UK | WT | 0.00 (0.04) vs. 0.00 (0.00) | 35.49 vs. 32.27 | 0.330 |
|  |  | MT | 0.00 (0.00) vs. 0.00 (0.01) | 35.99 vs. 38.93 | 0.413 |
|  | USA | WT | 0.00 (0.21) vs. 0.00 (0.00) | 36.51 vs. 27.66 | 0.015* |
|  |  | MT | 0.00 (0.04) vs. 0.00 (0.09) | 33.10 vs. 34.98 | 0.619 |
| Orf6 nonsynonymous | UK | WT | 0.00 (0.00) vs. 0.00 (0.00) | 36.42 vs. 31.19 | 0.052 |
|  |  | MT | 0.00 (0.08) vs. 0.02 (0.11) | 34.96 vs. 39.91 | 0.277 |
|  | USA | WT | 0.00 (0.15) vs. 0.00 (0.00) | 34.26 vs. 30.38 | 0.286 |
|  |  | MT | 0.00 (0.00) vs. 0.00 (0.00) | 36.03 vs. 31.78 | 0.133 |
| Orf7a synonymous | UK | WT | 0.00 (0.10) vs. 0.00 (0.00) | 37.36 vs. 30.10 | 0.076 |
|  |  | MT | 0.00 (0.00) vs. 0.00 (0.05) | 31.47 vs. 43.21 | 0.005* |
|  | USA | WT | 0.00 (0.05) vs. 0.00 (0.00) | 36.94 vs. 27.14 | 0.003* |
|  |  | MT | 0.00 (0.03) vs. 0.00 (0.02) | 34.83 vs. 33.09 | 0.647 |
| Orf7a nonsynonymous | UK | WT | 0.00 (0.06) vs. 0.00 (0.15) | 32.69 vs. 35.52 | 0.485 |
|  |  | MT | 0.00 (0.06) vs. 0.08 (0.11) | 28.51 vs. 46.01 | <0.001* |
|  | USA | WT | 0.05 (0.12) vs. 0.00 (0.00) | 36.89 vs. 27.21 | 0.021* |
|  |  | MT | 0.00 (0.03) vs. 0.00 (0.07) | 32.59 vs. 35.55 | 0.463 |
| Orf7b synonymous | UK | WT | 0.00 (0.00) vs. 0.00 (0.00) | 36.00 vs. 31.68 | 0.088 |
|  |  | MT | 0.00 (0.00) vs. 0.00 (0.00) | 35.50 vs. 39.39 | 0.047* |
|  | USA | WT | 0.00 (0.00) vs. 0.00 (0.00) | 32.27 vs. 32.78 | 0.797 |
|  |  | MT | 0.00 (0.07) vs. 0.00 (0.39) | 32.46 vs. 35.69 | 0.410 |
| Orf7b nonsynonymous | UK | WT | 0.00 (0.00) vs. 0.00 (0.00) | 35.53 vs. 32.23 | 0.219 |
|  |  | MT | 0.00 (0.00) vs. 0.00 (0.12) | 30.65 vs. 43.99 | 0.001* |
|  | USA | WT | 0.00 (0.00) vs. 0.00 (0.00) | 33.04 vs. 31.84 | 0.612 |
|  |  | MT | 0.00 (0.00) vs. 0.00 (0.00) | 32.27 vs. 35.89 | 0.153 |
| Orf8 synonymous | UK | WT | 0.00 (0.00) vs. 0.00 (0.00) | 37.68 vs. 33.21 | 0.534 |
|  |  | MT | 0.00 (0.00) vs. 0.00 (0.01) | 34.64 vs. 40.21 | 0.093 |
|  | USA | WT | 0.00 (0.00) vs. 0.00 (0.00) | 32.86 vs. 32.07 | 0.769 |
|  |  | MT | 0.00 (0.00) vs. 0.00 (0.00) | 34.43 vs. 33.53 | 0.791 |
| Orf8 nonsynonymous | UK | WT | 0.08 (0.23) vs. 0.00 (0.46) | 31.75 vs. 36.61 | 0.278 |
|  |  | MT | 0.00 (0.05) vs. 0.09 (0.12) | 25.79 vs. 48.59 | <0.001* |
|  | USA | WT | 2.46 (0.37) vs. 1.64 (0.95) | 43.40 vs. 19.34 | <0.001* |
|  |  | MT | 0.25 (0.47) vs. 0.22 (0.48) | 34.04 vs. 33.95 | 0.985 |
| Orf10 synonymous | UK | WT | 0.00 (0.00) vs. 0.00 (0.00) | 33.96 vs. 34.06 | 0.932 |
|  |  | MT | 0.00 (0.00) vs. 0.00 (0.00) | 34.94 vs. 39.92 | 0.050* |
|  | USA | WT | 0.00 (0.00) vs. 0.00 (0.00) | 32.00 vs. 33.10 | 0.272 |
|  |  | MT | 0.00 (0.05) vs. 0.00 (0.00) | 34.97 vs. 32.94 | 0.548 |
| Orf10 nonsynonymous | UK | WT | 0.00 (0.13) vs. 0.00 (0.00) | 36.54 vs. 31.05 | 0.140 |
|  |  | MT | 0.00 (0.07) vs. 0.00 (0.13) | 36.19 vs. 38.74 | 0.541 |
|  | USA | WT | 0.00 (0.22) vs. 0.00 (0.00) | 36.80 vs. 27.31 | 0.004* |
|  |  | MT | 0.00 (0.00) vs. 0.00 (0.00) | 35.27 vs. 32.61 | 0.220 |

IQR, Interquartile Range; * Wilcoxon rank-sum (Mann-Whitney) test; *p-value<0.05 is significant.
